# Characterizing enteric neurons in Dopamine Transporter (DAT)-Cre reporter mice reveals dopaminergic subtypes with dual-transmitter content

**DOI:** 10.1101/2023.06.16.545271

**Authors:** Sherilyn Junelle Recinto, Shobina Premachandran, iparna Mukherjee, Alexis Allot, Adam MacDonald, Moein Yaqubi, Samantha Gruenheid, Louis-Eric Trudeau, Jo Anne Stratton

## Abstract

The enteric nervous system (ENS) comprises a complex network of neurons whereby a subset appears to be dopaminergic, although the characteristics, roles, and implications in disease are less understood. Most investigations relating to enteric dopamine (DA) neurons rely on immunoreactivity to tyrosine hydroxylase (TH) - a rate-limiting enzyme in the production of DA. However, TH immunoreactivity is likely to provide an incomplete picture given previous work has showed that some DA neurons contain little if any TH and its levels tend to be decreased in response to cellular stress. This study herein provides a comprehensive characterization of DA neurons in the gut using a well-accepted reporter mouse line, expressing a fluorescent protein (tdTomato) under control of the DA transporter (DAT) promoter. Our findings confirm a unique localization of DA neurons in the gut and unveil the discrete subtypes of DA neurons in this organ, which we characterized using both immunofluorescence and single-cell transcriptomics, as well as validated using *in situ* hybridization. We observed distinct subtypes of DAT-tdTomato neurons expressing co-transmitters and modulators across both plexuses; some of them likely co-releasing acetylcholine, while others were positive for a slew of canonical DA markers (TH, VMAT2 and GIRK2). Interestingly, we uncovered a seemingly novel population of DA neurons unique to the ENS which were ChAT/DAT-tdTomato-immunoreactive neurons and were characterised by the expression of *Grp*, *Calcb* and *Sst*. Given the clear heterogeneity of DAergic gut neurons, further investigation is warranted to define their functional signatures and discover any inherent vulnerabilities in disease.

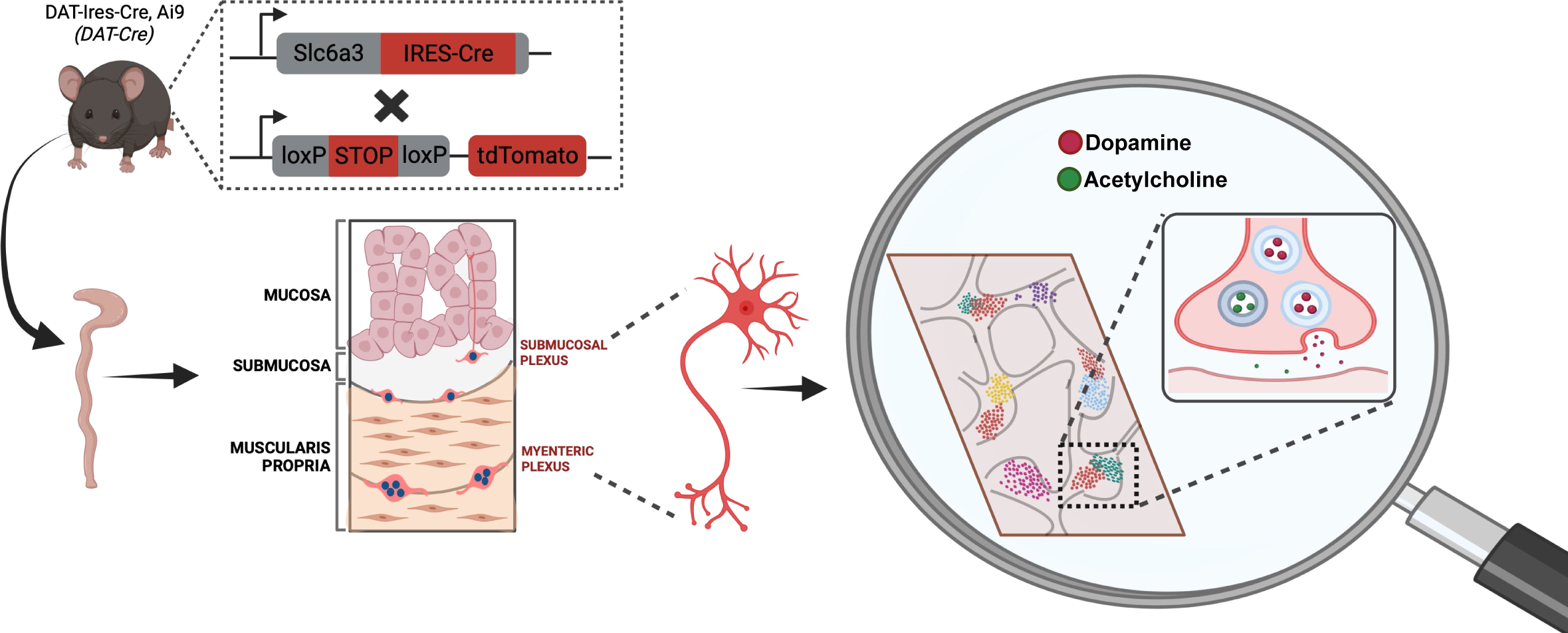

Using a reporter mouse line, expressing a fluorescent protein under control of the dopamine transporter (DAT) promoter, discrete subtypes of dopaminergic neurons were unveiled across the ganglionated plexuses of the gut. A novel subpopulations of enteric DA neurons, expressing genes previously reported involved in dopamine signaling in the brain, exhibit a cholinergic phenotype.

## Introduction

An altered gut-brain axis is increasingly recognized as central to many neurological conditions including in Parkinson’s disease (PD). PD is the second most prevalent neurodegenerative disease, affecting more than 10 million people globally (Ou et al., 2021). Its pathological characteristics are commonly defined by the accumulation of Lewy body inclusions and by the gradual degeneration of dopamine (DA) neurons in the substantia nigra pars compacta (SNc) of the midbrain. The selective loss of DA is thought to underlie some of the canonical motor symptoms of PD (Greffard et al., 2006). There is growing recognition that non-motor abnormalities, including perturbed sleep, anosmia, depression and gut dysmotility precede motor deficits by up to 20 years (Gaenslen et al., 2011; Pont-Sunyer et al., 2015; Schrag et al., 2015). These are likely to result from the dysfunction or loss of multiple types of neurons, albeit quantitative evidence for the extent of neuronal loss in regions other than the SNc is presently fragmented (Giguère et al., 2018). Although it is likely that most PD patients suffer from some level of dysfunction of the peripheral nervous system (PNS) (Abbott et al., 2001; Borghammer, 2023; Fedorova et al., 2017; Raeder et al., 2023; Savica et al., 2009; Skjærbæk et al., 2021), whether gut dysmotility and associated constipation are driven in part by loss of enteric DA neurons remains largely unexplored (Anderson et al., 2007; Francesca et al., 2023; Liu et al., 2021; Wang et al., 2012). Various histological studies have demonstrated the presence of Lewy body inclusions in areas outside the central nervous system (CNS) predominantly in the gastrointestinal tract (Del Tredici & Braak, 2012; Dickson et al., 2009). These inclusions can be found in neurons residing in the gut and positively correlate with constipation in PD patients (Lebouvier et al., 2010). Pathological features observed from colonic biopsies were in fact apparent at least 8 years prior to PD diagnosis (Hilton et al., 2014). Further reports revealed neuronal death in the ganglionated plexus of patients’ colons as well as in rodent models of PD (Lebouvier et al., 2010; McQuade et al., 2021; Ohlsson & Englund, 2019). Despite clear involvement of the gastrointestinal tract in PD, there are inconsistencies reported for the extent of neuronal abrogation (O’Day et al., 2022). The discrepancies in present studies may be largely due to the heterogeneity in clinical presentation and data collection; thus, a balanced criteria for patient recruitment, consistency in neuronal quantification methods and study design, as well as objective evaluations of constipation will be instrumental in teasing apart the scope of enteric neurodegeneration in PD. Furthermore, adding dedicated analysis for studying neuronal subtypes affected in the gut of PD patients will also be critical.

Recent single-cell transcriptomics studies in both human and mice have started to explore the complexity of the cellular landscape within the gut including defining unique gene expression signatures within the neuronal lineage (Drokhlyansky et al., 2020; May-Zhang et al., 2021; Memic et al., 2018; Morarach et al., 2021; Wright et al., 2021). However, very few of these studies have included the identification of catecholaminergic neuronal cell types expressing genes involved in DA synthesis, such as tyrosine hydroxylase (*TH*) (Memic et al., 2018; Morarach et al., 2021; Wright et al., 2021). Amongst these studies, only one has explored this population describing novel markers in the developing ENS of fetal mice, but this may not be translatable to the expression patterns of mature DA neurons (Memic et al., 2018). As such, rigorous studies describing the diversity of DA neuronal populations in the gut have not yet been carried out. Historically, a neuronal subtype is defined by the primary neurotransmitter it releases and its functional properties. With the gradual appreciation that many if not most neuronal types in the CNS and PNS utilize more than one type of neurotransmitter (El Mestikawy et al., 2011; Trudeau, 2004; Trudeau & El Mestikawy, 2018; Trudeau & Gutiérrez, 2007) and with the advent of single-cell transcriptomic approaches, assigning a molecular gene signature is progressively being used to precisely define neuronal sub-classes. In the case of midbrain DA neurons, previous microarray and recent single-cell RNA sequencing (RNAseq) work has revealed extensive heterogeneity including in genes specifying their developmental origin, as well as a battery of genes involved in the synthesis and vesicular packaging of DA, but also of other neurotransmitters including glutamate and various neuropeptides (Gaertner et al., 2022; Kouwenhoven et al., 2020; Poulin et al., 2018; Poulin et al., 2020; Poulin et al., 2014).

The enteric nervous system (ENS) is a network of neurons situated in two separate regions of the gastrointestinal wall; the submucosal plexus (SMP) found underneath the mucosa and the myenteric plexus (MP) located within the muscularis propria. The two plexuses are responsible for varying functions of the gut; the SMP plays a crucial role in secretion and luminal sensing, whereas the MP regulates gastrointestinal motility through peristaltic waves. These complex intestinal functions require a diverse population of neurons including TH-positive DA neurons as well as serotonin, cholinergic and nitrergic neurons (Furness et al., 2014). Although there are several reports of the presence of DA neurons in the gut (Li et al., 2004; Walker et al., 2000), such studies typically rely solely on immunostaining for TH – the rate limiting enzyme in the synthesis of catecholamines to define a DAergic phenotype. Further, DA neuron subtyping analysis has only been characterised, to some extent, in the murine ileum, demonstrating that some TH neurons are immunoreactive for the DA transporter (DAT) (and others are not), as well as vice-versa (Li et al., 2004).

The use of transgenic reporter mouse lines such as those in which promoters for *DAT* drive Cre recombinase to induce the expression of fluorescent molecules, such as tdTomato (named here as DAT-Cre mice) has the potential to shed new light on enteric DA neuron diversity. By taking advantage of DAT-Cre mice, we sought to deepen our understanding of DA neurons and their putative subtypes in the ENS. We conducted immunofluorescence staining in colonic tissues along with single-cell RNAseq and *in situ* hybridization to unravel the heterogeneity of DAergic neuronal populations. Providing an in-depth characterization of these classes of enteric neurons will be conducive to future endeavors aimed at better understanding their functional signatures and any potential selective vulnerabilities of enteric neurons in disease.

## Methods

Protocols associated with this work can be found on protocols.io: dx.doi.org/10.17504/protocols.io.3byl4q792vo5/v1

### Mice and tissue processing

*DAT-Ires-Cre* (RRID:IMSR_JAX:006660) mice were bred with Ai9 mice (RRID: IMSR_JAX:007909) to induce the expression of tdTomato in DAT-expressing cells (DAT-Cre mice). Adult DAT-Cre mice (both sexes) of age 2-3 months were utilized in the present study. Mice were housed under pathogen-free conditions, given water and food *ad libitum.* Studies were approved by the Université de Montréal Animal Care and Use Committee (CDEA). CDEA guidelines on the ethical use and care of animals were followed. The mice were euthanized by isoflurane followed by CO_2_ asphyxiation. Age-matched wild type C57Bl6/J mice (RRID:IMSR_JAX:000664) were also used as control for tdTomato expression. We collected gut tissue, and focused on colons where fewer investigations of the ENS have been made. After isolation, fecal material was flushed with an 18-gauge needle and adipose tissue was removed. Colons were cut open longitudinally and coiled around themselves, resembling a *swiss-roll* (as depicted in Fig 1A), then fixed in 4% paraformaldehyde (PFA) for 24 hours and dehydrated with a 30% sucrose solution. The tissue was then embedded and cryosectioned serially to obtain coronal sections at 15 μm, then stored at -80°C. Of note, we used a *swiss-roll* preparations to easily compare dopaminergic neurons between ganglionated plexuses within the same tissue section. This is of importance as previous studies have underlined the varying density of TH-immunoreactive neurons within the ENS (Chalazonitis et al., 2022; Li et al., 2004; Qu et al., 2008), which is pertinent given that specific locations of neuronal populations may influence their functions. It is also noted in disease, such as PD, that submucosal neurons appear to be selectively vulnerable (Francesca et al., 2023; Lebouvier et al., 2010; Stokholm et al., 2016).

**Figure 1:**
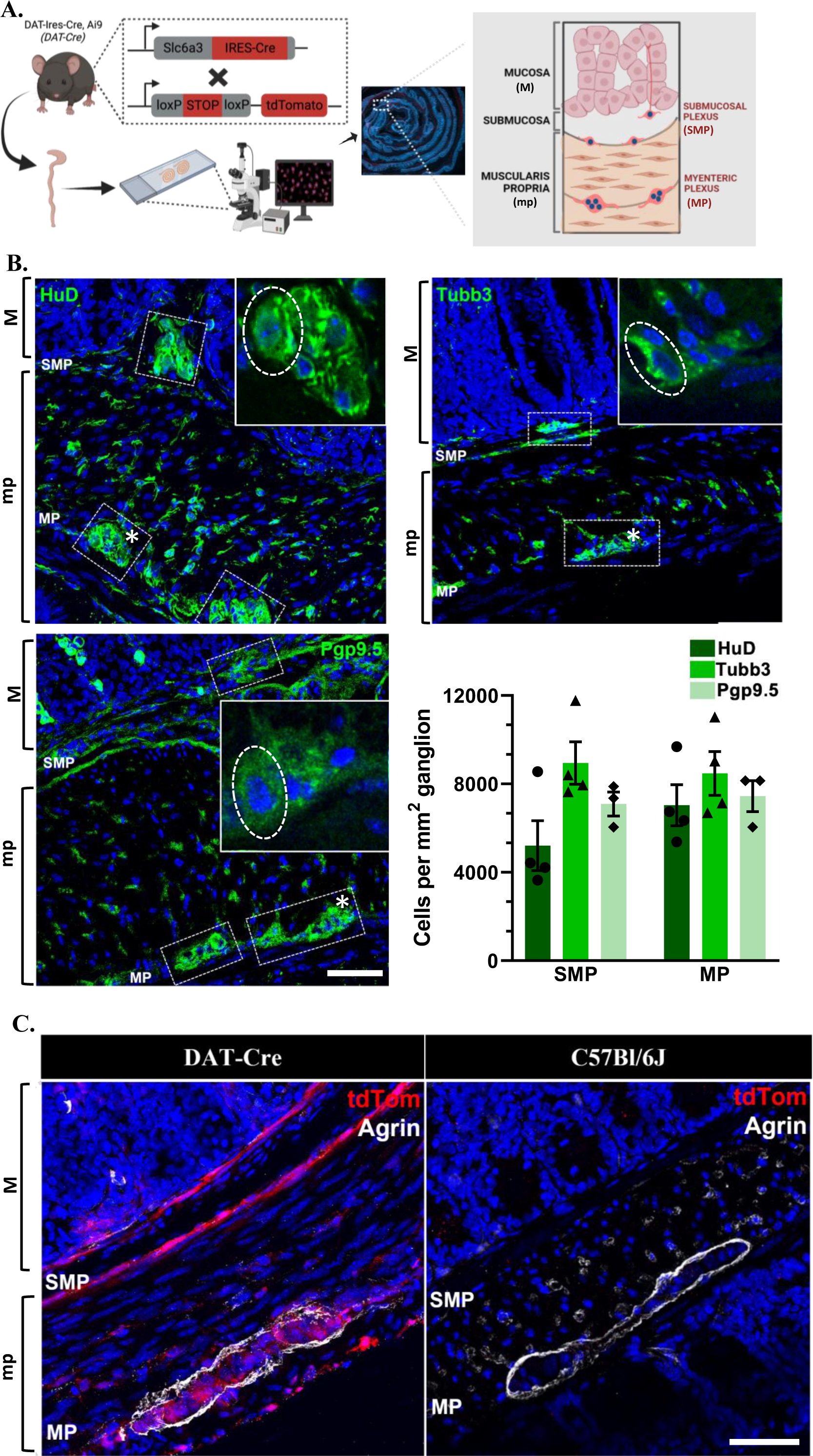
Detection of enteric neurons in DAT-tdTomato transgenic mice. **A**) Schematic diagram of immunofluorescence staining protocol for mouse colon isolated from transgenic mice, where tdTomato (Ai9) is controlled under the promoter for dopamine transporter (Slc6a3, referred to as DAT-Cre). The coronal plane of the colon mounted in a *“swiss-roll”* depicts the three subdivision of intestinal layers and the two ganglionated plexuses. The submucosal plexus (SMP) is shown between the mucosa (M) and the muscularis propria (mp), while the myenteric plexus (MP) is within the mp. **B)** Representative immunofluorescence staining with cross-sectional view of colonic tissue from DAT-Cre mice. Enteric neurons within the two ganglionated plexuses are detectable with several pan-neuronal markers that are commonly used to identify neurons. Neurons (green) are immunoreactive for RNA-binding human antigen (HuD), tubulin β3 (Tubb3) and ubiquitin carboxyl-terminal hydrolase-1 (Uch-l1/Pgp9.5). Ganglia are outlined by white dotted box. The inset is a magnified view of the ganglion designated with * and an individual cell is delineated by white dotted lines. The density of neurons normalized to ganglionic area (in mm^2^) is quantified in both plexuses. Mean ± SEM, n=4 mice per group with 3 ganglia quantified per mouse. **C)** Cross-sectional view of colonic tissue from DAT-Cre mice immunoreactive for tdTomato (red). Agrin is used as a marker for blood-myenteric-barrier (white) to depict ganglia. Note the lack of tdTomato signal in wild type C57Bl/6J colonic tissue, as expected. Nuclei are stained with Hoechst (blue). Scale bars: 50 μm.

### Immunofluorescence

Following thawing, tissue sections were permeabilized and blocked (10% donkey serum with 0.5% Triton X-100) for 2h. Primary antibody stains were performed overnight at room temperature (RT) followed by secondary antibody staining for 2h. Primary and secondary antibodies used are indicated in **Table 1**. Hoechst 33258 was used for nuclear staining (1:5000, ThermoFisher H3569, RRID:AB_2651133). Slides were mounted in Permafluor Mounting media (ThermoFisher TA030FM).

**Table 1:** Primary antibodies used in immunofluorescence staining of colonic tissue.

| Antigen | Host | Dilution | Company (CAT#) | RRDI |
| --- | --- | --- | --- | --- |
| Tubb3 | Mouse | 1:500 | Millipore (MAB5564) | RRID:AB_570921 |
| Pgp9.5 | Mouse | 1:500 | ThermoFisher (480012) | RRID:AB_2532243 |
| HuD | Rabbit | 1:200 | Abcam (ab96474) | N/A |
| ChAT | Goat | 1:200 | Sigma Aldrich (SAB2500236) | RRID:AB_10603616 |
| nNOS | Rabbit | 1:500 | ThermoFisher (61-7000) | RRID:AB_2313734 |
| tdTomato | Goat | 1:200 | Rockland (200-101-379) | RRID:AB_2744552 |
| tdTomato | Rabbit | 1:200 | Rockland (600-401-379) | RRID:AB_2209751 |
| Th | Chicken | 1:100 | Millipore (AB9702) | RRID:AB_570923 |
| Girk2 | Rabbit | 1:100 | Alomone Labs (APC-006) | RRID:AB_2040115 |
| Foxa2 | Rabbit | 1:100 | R&D Systems (AF2400) | RRID:AB_2294104 |
| Vmat2 | Rabbit | 1:200 | Abcam (ab70808) | RRID:AB_1952699 |
| Anti goat AF488 | Donkey | 1:200 | Thermo Fisher (A11055) | RRID:AB_2534102 |
| Anti rabbit AF488 | Donkey | 1:200 | Thermo Fisher (A-21206) | RRID:AB_2535792 |
| Anti goat AF555 | Donkey | 1:200 | Thermo Fisher (A-21432) | RRID:AB_2535853 |
| Anti mouse AF555 | Donkey | 1:200 | Thermo Fisher (A-31570) | RRID:AB_2536180 |
| Anti rabbit AF555 | Donkey | 1:200 | Thermo Fisher (A-31572) | RRID:AB_162543 |
| Anti mouse AF647 | Donkey | 1:200 | Thermo Fisher (A-31571) | RRID:AB_162542 |
| Anti chicken AF647 | Donkey | 1:200 | Jackson ImmunoResearch (703-605-155) | RRID:AB_2340379 |

### *In situ* hybridization

Following thawing, tissue sections were heated for 30 mins at 60°C, post-fixed in pre-chilled 4% PFA for 15 mins at 4°C, and dehydrated by subsequent immersion in 50, 70, and 100% ethanol for 5 mins at RT. *In situ* staining of fixed-frozen samples were performed according to the RNAscope Multiplex Fluorescence manufacturer’s instructions (Advanced Cell Diagnostics, ACD). Briefly, tissue sections were incubated for 10 mins at 95°C for antigen retrieval and Protease III was applied for 30 mins at 40°C. After multiple washes, RNAscope multiplex assay was carried out as described in ACD’s protocol followed by the aforementioned immunofluorescence protocol to co-stain with tdTomato and ChAT antibodies. All incubations were at 40°C and used a humidity control chamber. Mouse probes used were Grp-C1, Calcb-C1 and Sst-C1; and Opal dye 520 was used at 1:1500 concentration. Hoechst 33258 was used for nuclear staining (1:5000, ThermoFisher H3569). Slides were mounted in Prolong Gold Antifade Mounting media (ThermoFisher P36930).

### Single-cell suspension preparation

Given protocols are established for longitudinal muscle and myenteric plexus (LMMP) isolations, we prepared single cell suspensions from this intestinal layer. Isolated colons from adult DAT-Cre mice (n=4, equal sexes) were washed, cut open longitudinally as described above, and pinned with the intestinal lumen facing down on a 4% agarose pad. The LMMP was then micro-dissected by peeling it off using fine-tip tweezers, and was then collected and cut into 1-2 cm pieces. The tissues were then enzymatically dissociated to release myenteric neurons and other cell types using 1 mg/mL collagenase IV (Sigma, C5138) and 0.05% trypsin (Sigma, T4049), adapted from (Smith et al., 2013). Dissociated cells were filtered through a 40 μm cell strainer and stained with eFluor506 fixable viability dye (ThermoFisher 65-0866-14). Using fluorescence-activated cell sorting (FACS Aria), we detected ∼60% of events were live single cells and of that ∼20% were DAT-tdTomato-positive.

### Single-cell library preparation and analysis

Twenty thousand live cells were FACS-collected and then prepared for single-cell sequencing. All cells were processed according to 10X Genomics Chromium Single Cell 3’ Reagent Guidelines (https://support.10xgenomics.com/single-cell-gene-expression). Briefly, cells were partitioned into nanoliter-scale Gel Bead-In-EMulsions (GEMs) using 10X GemCode Technology. Primers containing (i) an Illumina R1 sequence, (ii) a 16 bp 10x barcode, (iii) a 10 bp Unique Molecular Identifier (UMI) and (iv) a poly-dT primer sequence were incubated with partitioned cells resulting in barcoded, full-length cDNA from poly-adenylated mRNA. Silane magnetic beads were used to remove leftover biochemical reagents/primers, then cDNA was amplified by PCR. Enzymatic fragmentation and size selection was used to optimize cDNA amplicon size prior to library construction. R1 (read 1 primer sequence) were added during GEM incubation, whereas P5, P7, a sample index (i7), and R2 (read 2 primer sequence) were added during library construction via end repair, A-tailing, adaptor ligation and PCR. Quality control and quantification was performed using an Agilent Bioanalyzer High Sensitivity chip. Sequencing was performed using NovaSeq 6000 S4 PE 100bp, which resulted in a read depth of 46 000 reads/cell. Reads were processed using the 10X Genomics Cell Ranger Single Cell 2.0.0 pipeline (RRID:SCR_017344, https://support.10xgenomics.com/single-cell-gene-expression/software/pipelines/latest/what-is-cell-ranger) with default and recommended parameters, as previously described (Zheng et al., 2017). FASTQs generated from sequencing output were aligned to the mouse GRCm38.p5 (primary assembly) reference genome using the STAR algorithm 2.7.3a (RRID:SCR_004463, http://code.google.com/p/rna-star/) (Rahmani et al., 2020). Next, gene-barcode matrices were generated for each individual sample by counting unique molecular identifiers (UMIs) and filtering non-cell associated barcodes. This output was then imported into the Seurat v4.9.9.9039 R toolkit v.4.2.1 (RRID:SCR_016341, https://satijalab.org/seurat/get_started.html; RRID:SCR_001905, http://www.r-project.org/) for quality control and downstream analysis of our single-cell RNA-seq experiment (Macosko et al., 2015). All functions were run with default parameters, unless specified otherwise. Low quality cells (<200 genes/cell and >5 % of mitochondrial genes) were excluded from the overall experiment. Gene expression was log normalized to a scale factor of 10 000. A total of ∼8, 000 cells and ∼20, 000 genes were detected.

### Tissue quantification and statistical analysis

Immunostained tissue slices were imaged using a Leica SP8 Confocal microscope and quantified with the Las X Leica software. Images were taken at 40X magnification with Z-stacks (step size of 0.9). Each ganglion was identified in either the submucosal or myenteric plexus based on morphology as shown in earlier publications (Kulkarni et al., 2018; Li et al., 2004). The quantification was unbiasedly performed using Z-stacks where the assessor defined a nucleus within the ganglia, then scroll through the Z-stack to ensure that the stains were consistently expressed in and around the cell body surrounding the nuclei. Tubb3 and tdTomato (as well as most of the other stains used) have clear cytoplasmic signal abutting the nuclei, which made this approach ideal for ensuring accurate quantification. All quantitative analysis at the ganglia was normalized to an area of 900 μm^2^ (calculated as the average size of a ganglion). The number of animals used with 3-4 ganglia analyzed per colon at each plexus can be found in the figure legends. All statistical analysis was performed using GraphPad Prism 5 (RRID:SCR_002798, http://www.graphpad.com/). For comparisons across multiple groups, a two-way analysis of variance (ANOVA) followed by Tukey’s post-hoc comparison was used. For comparisons of across two groups, an unpaired Student’s t test was used. P values of < 0.05 were considered significant.

For quantification of single-cell RNAseq data, unsupervised clustering of cells was performed. A selection of highly variable genes (2000 genes) was obtained and used for principal component analysis. Significant principal components were determined using JackStraw analysis. PCs 1 to 20 were used for graph-based clustering (at resolution = 1) to identify distinct groups of cells. These groups were projected onto UMAP analysis using previously computed principal components 1 to 20. Expression of selected marker genes (*Elavl4*, *Tubb3, Uchl1* and *Map2*) was used to classify neurons and these initial clusters were further divided into seven cell clusters identified by graph-based clustering.

## Results

### Enteric pan-neuronal markers and tdTomato expression are detectable in both colonic plexuses in DAT-tdTomato transgenic mice

DAT-Cre transgenic lines are considered more reliable in the selective identification of midbrain DA neurons compared to other reporter mouse lines such as TH-Cre (Bäckman et al., 2006; Ekstrand et al., 2007; Engblom et al., 2008; Lindeberg et al., 2004; Zhuang et al., 2005). As such, we used DAT-Cre mice crossed with Ai9 cre-dependent tdTomato mice and immunohistochemistry (Fig. 1A) to confirm the existence of submucosal and myenteric DA neurons, as well as characterize their heterogeneity. Using cross sectional analysis in *swiss-roll* format, we detected three subdivisions of the intestinal layers and two ganglionated plexuses, as expected. We consistently detected the submucosal plexus (SMP) between the mucosa and the muscularis propria, as well as the myenteric plexus (MP) within the muscularis propria, using a battery of pan-neuronal markers (Fig. 1B) (RNA-binding human antigen, HuD; tubulin β3, Tubb3; and ubiquitin carboxyl-terminal hydrolase-1, Uch-l1/Pgp9.5). We explored the efficacy of these commonly used pan-neuronal markers given, in most cases, they are utilized interchangeably, with little evidence to support which markers are most robust. Tubb3 is a cytoskeletal protein involved in microtubule stability and is often used to label both neuronal soma and axonal projections in the CNS or PNS, whereas HuD and Pgp9.5 are expressed predominantly in peripheral neurons (Bolognani et al., 2010; Hibi et al., 1999; Liu et al., 2007). A direct comparison of the expression of these neuronal markers revealed that there was no statistical difference within or between plexuses across all markers (Fig. 1B). We opted to focus largely on Tubb3 as a co-label with tdTomato for consistency and due to compatibility considerations with other antibodies. Of note, we identified non-neuronal tdTomato-positive signal in the vicinity of the plexuses (data not shown) which are likely immune cells (Gopinath et al., 2022; Mackie et al., 2018), and thus we were rigorous in performing all analysis with pan-neuronal marker co-labeling. We also determined that tdTomato expression was robust in colonic tissue in both the SMP and MP of DAT-Cre mice, but that the signal was not detected in wild type C57Bl/6J colonic tissue, as expected (Fig. 1C). We used a blood-myenteric-barrier marker to define MP borders, termed Agrin (which is not expressed in the SMP) (Dora et al., 2021), important for defining ganglia localization in the MP (Fig. 1C).

### TdTomato-immunoreactive DA neurons vary across plexuses

In keeping with previous descriptions of DA neurons in the gut demonstrating that the density of enteric TH-expressing neurons gradually decreases from rostral to caudal region of the gastrointestinal tract and from the SMP to MP (Chalazonitis et al., 2022; Li et al., 2004; Qu et al., 2008), we found a reduced density of tdTomato/Tubb3-positive DA neurons in the MP compared to the SMP using DAT-Cre mice (Fig. 2A). To ascertain the DAergic phenotype of tdTomato-positive neurons, we also co-label these cells with TH, the rate-limiting DA synthesis enzyme. We indeed observed that most of these neurons were also immunoreactive for TH in ganglionated plexuses. Interestingly, a small proportion of neurons were positive for TH but not for tdTomato, consistent to what has been described in the ventral midbrain (Lammel et al., 2015; Poulin et al., 2018; Tiklová et al., 2019). By contrast, we noted few submucosal and myenteric ganglia containing tdTomato-immunoreactive neurons but negative for TH (Fig. 2B).

**Figure 2:**
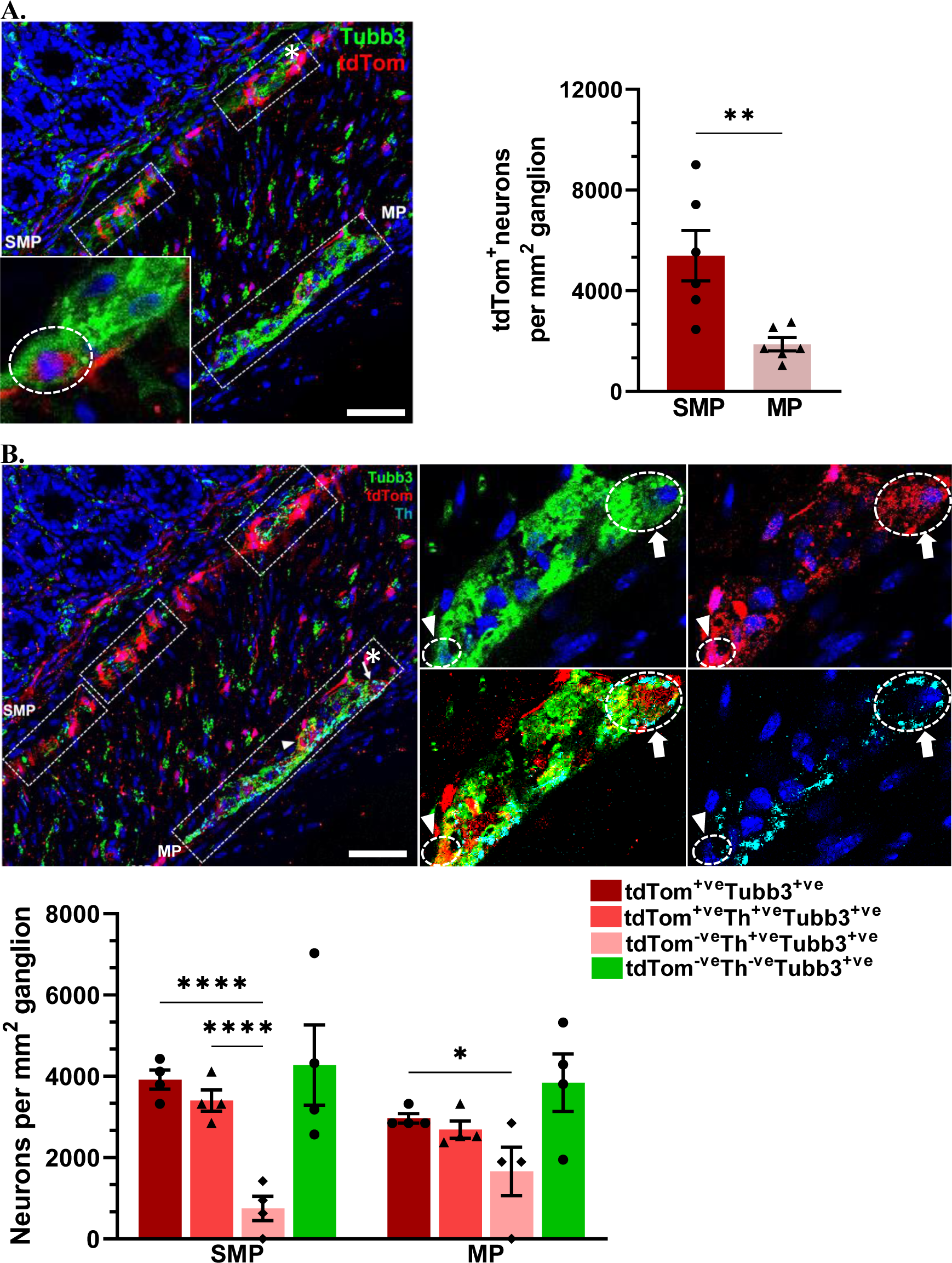
TdTomato-immunoreactivity varies across plexuses and colocalizes with tubulin β3. **(Tubb3) and tyrosine hydroxylase (Th)-positive cells. A)** Representative coronal slice of colonic tissue illustrating that DAT-tdTomato -expressing cells (red) are immunoreactive for a pan-neuronal marker, Tubb3 (green) in the submucosal and myenteric plexuses (SMP, MP). The density of Tubb3^+ve^tdTom^+ve^ neurons normalized to ganglionic area (in mm^2^) is quantified in both plexuses, which revealed significantly less DAT-tdTomato^+ve^ neurons in the MP compared to the SMP. Unpaired, two-tailed t-test, p<0.05. Mean ± SEM, n=6 mice per group with 3 ganglia per mouse quantified. **B)** Representative immunofluorescence staining for tyrosine hydroxylase (Th) (cyan), which is a rate limiting enzyme for generating L-3,4-dihydroxyphenylalanine (L-DOPA), the precursor for dopamine. Th^+ve^ cells colocalized with tdTomato (red) and Tubb3 (green) in both ganglionated plexuses. The density of Tubb3^+ve^ enteric neurons expressing either: 1) DAT-tdTomato (tdTom^+ve^Tubb3^+ve^); 2) DAT-tdTomato and Th (tdTom^+ve^Th^+ ve^Tubb3^+ve^); 3) Th^+ve^ alone (tdTom^-ve^Th^+ve^Tubb3^+ve^); or 4) No dopamine neuron markers (tdTom^-ve^Th^-ve^Tubb3^+ve^); is presented as normalized to ganglionic area (in mm^2^). This revealed that most Th-expressing neurons in the gut co-localize with DAT-tdTomato in both plexuses but a smaller subset do not express DAT-tdTomato. Two-way ANOVA followed by Tukey’s multiple comparison’s test, p<0.05. Mean ± SEM, n=4 mice per group with 3 ganglia per mouse quantified. Ganglia are outlined by the white dotted box. The inset is a magnified view of the ganglion designated with * and an individual cell is delineated by white dotted lines. Nuclei are stained in blue. Scale bars: 50 μm.

### A subpopulation of tdTomato-immunoreactive neurons in the ENS exhibit dual-transmitter content

The complex neural networks in the ENS can be largely categorized as either excitatory or inhibitory motor neurons (Brehmer, 2021). The former mostly express choline acetyltransferase (ChAT), generally classified as cholinergic, whereas the latter is commonly referred to as nitrergic due to their release of nitric oxide, an inhibitory molecule synthesized by neuronal-specific nitric oxide synthase (nNOS) (Dharshika & Gulbransen, 2023). The striking abundance of tdTomato/TH-positive neurons in the colon wall in the present study, especially in the SMP (Fig. 2A), led us to speculate that a subset of enteric cholinergic or nitrergic neurons may also have a DAergic phenotype. This conclusion would coincide with a growing literature demonstrating that DA neurons as well as other CNS and PNS neurons use a combination of neurotransmitters (Brunet Avalos & Sprecher, 2021; Dal Bo et al., 2004; El Mestikawy et al., 2011; Trudeau & El Mestikawy, 2018). To examine this, we performed co-immunolabelling for HuD and ChAT, Tubb3 and nNOS, tdTomato and ChAT, as well as tdTomato and nNOS. Firstly, we confirmed the presence of both cholinergic and nitrergic neurons in both plexuses at similar levels (Fig. 3A). Then we assessed ChAT- and nNOS-positive neurons for DAT-tdTomato (Fig. 3B). Indeed, we observed the existence of a double-phenotype DA neuron population in both the SMP and MP, which accounts to ∼20% and ∼50% of tdTomato-immunoreactive neurons, respectively. Of note, we also found that the majority of tdTomato-immunoreactive neurons do not express both ChAT and nNOS in the SMP, but nearly half in the MP do.

**Figure 3:**
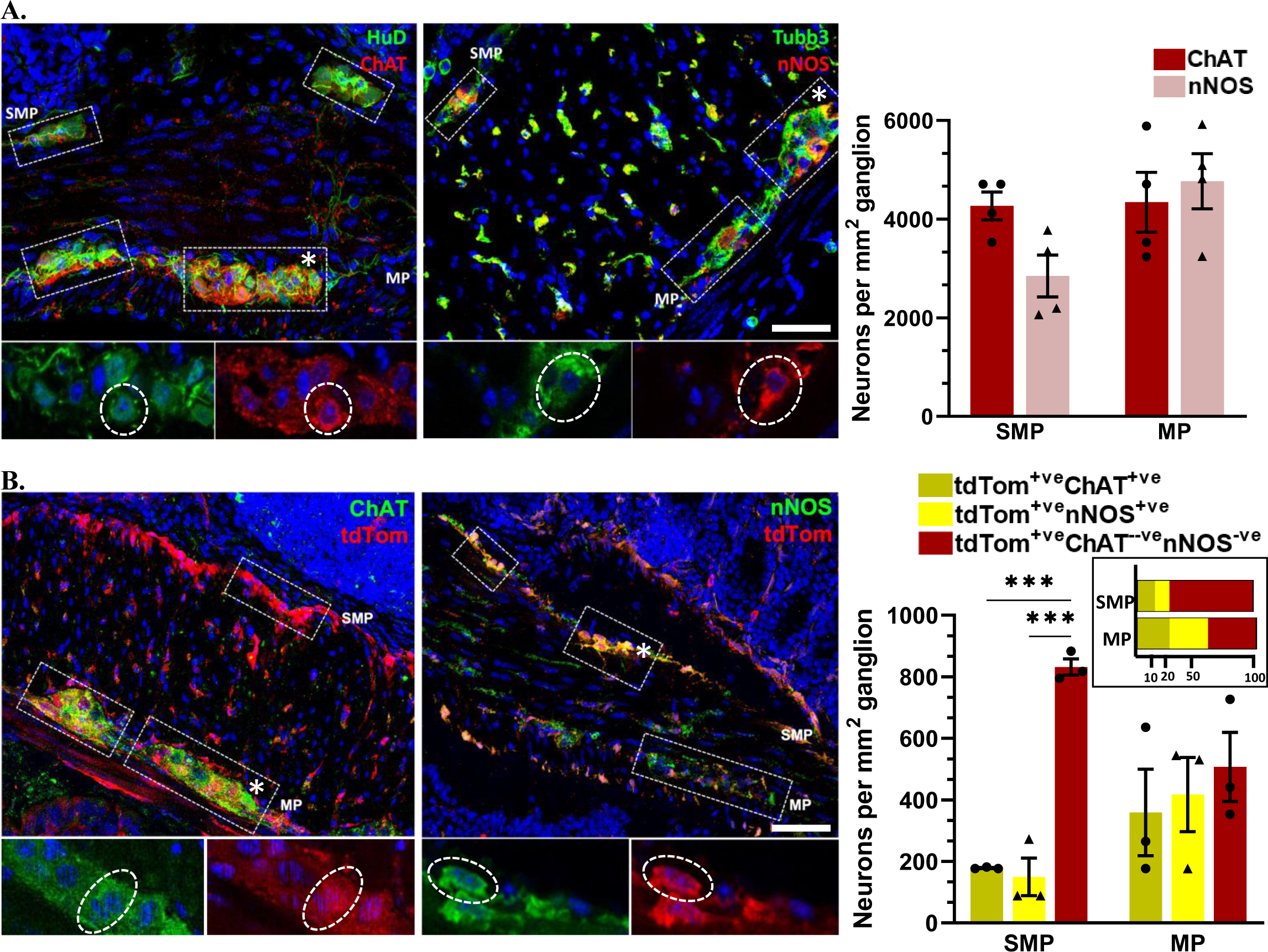
A subpopulation of tdTomato-immunoreactive neurons in the ENS exhibit dual-transmitter content. **A)** Representative cross-sectional view of colonic tissue labelling cholinergic (choline acetyl transferase, ChAT-positive) or nitrergic (neuronal nitric oxide synthase, nNOS-positive) (red) neurons (green, HuD or Tubb3) in submucosal and myenteric plexuses (SMP and MP). The density of ChAT^+ve^ or nNOS^+ve^ neuronal populations (colocalized with pan-neuronal markers HuD or Tubb3, respectively) are normalized to ganglionic area (in mm^2^). Mean ± SEM, n=4 mice per group with 3 ganglia per mouse quantified. **B)** Representative immunofluorescence staining for DAT-tdTomato-expressing cells (red) co-localizing with either ChAT or nNOS (green) in both ganglionated plexuses (SMP and MP). The densities of DAT-tdTomato neurons co-expressing either ChAT (tdTom^+ve^ChAT^+ve^) or nNOS (tdTom^+ve^nNOS^+ve^) are compared to that of DAT-tdTomato alone (tdTom^+ve^ChAT^-ve^nNOS^-ve^) in both plexuses normalized to ganglionic area (in mm^2^). The proportions of each subpopulations are shown in the inset as % relative to the total tdTomato^+ve^ cells. Two-way ANOVA followed by Tukey’s multiple comparison’s test, p<0.05. Mean ± SEM, n=3 mice per group with 3 ganglia per mouse quantified. Ganglia are outlined by white dotted box. The inset is a magnified view of the ganglion designated with * and an individual cell is delineated by white dotted lines. Nuclei are stained with Hoechst (blue). Scale bars: 50 μm.

### Single-cell transcriptomics corroborates the existence of ChAT-expressing dopaminergic neuron subtype

To further elucidate the molecular signature defining dopaminergic subtypes with dual-transmitter content, we used single-cell transcriptomics. Using four mice of equal sexes, we micro-dissected the LMMP from the colon followed by enzymatic digestion and FACS to obtain live single cells and generate a gene expression library (Fig. 4A). Following the application of computational quality control measures, we performed unsupervised clustering and annotated seven enteric clusters based on literature-curated cell type markers (Fig. 4B). Out of these cells, 247 were classified as neurons based on the expression of *Elavl4* (HuD), *Tubb3*, *Uchl1* (Pgp9.5) and *Map2*, which we further subdivided into seven distinct neuronal clusters (Fig. 4C; data also available at singlocell.openscience.mcgill.ca/EntericNeurons). Consistent with our immunostaining, we identified *Chat*-(Cluster 0, 2, 3 and 6) and *Nos1* (Cluster 1)-expressing neuronal cell clusters as the most prevalent cell types (Fig. 4C). We then evaluated *Th* expression and found highest expression in Clusters 2 and 5 (Fig. 5A). Notably, Cluster 5 showed higher expression of other canonical DA neuron markers, important in the vesicular packaging of DA (*i.e.,*vesicular monoamine transporter 2, *Vmat2*), in the regulation of SNc neurons excitability (*i.e.,* G-protein-regulated inward-rectifier potassium 2, *Girk2*), as well as previously reported genes expressed in midbrain DA neurons, such as vesicular glutamate transporter 2, *Vglut2*, cocaine- and amphetamine-regulated transcript protein, *Cartpt* and 5-hydroxytryptamine/serotonin receptor 2B, *Htr2b* (Ásgrímsdóttir & Arenas, 2020; Carpenter et al., 2020; Doly et al., 2017; El Mestikawy et al., 2011; Trudeau, 2004; Trudeau & El Mestikawy, 2018; Trudeau & Gutiérrez, 2007) (Fig. 5B). Validating our transcriptome observations, we detected immunoreactivity for VMAT2 and GIRK2 in DAT-tdTomato-expressing neurons within the ganglionated plexuses (Fig. 5C). As for Cluster 2, these *Th*-expressing neurons had very little to no expression of the aforementioned traditional DA neuron markers (Fig. 5B); however high *Chat* expression was noted in this cluster compared to *Th*-expressing Cluster 5 (Fig. 4C). We thus further explored the transcriptomic signature of this unique *Th/Chat*-expressing, DAergic neuronal subtype (Fig. 6A); and amongst them we detected a range of genes previously reported in the ENS (*Grp, Calcb* and *Sst*), but not yet shown to be present in enteric DA neurons (Fig. 6B). Interestingly, these genes have also been associated with DA signaling in the brain (Charbit et al., 2009; Elina et al., 2023; Kramer et al., 2018; Salesse et al., 2020). Validation experiment using *in situ* hybridization indicated the expression of these novel markers in a small fraction of cells immunoreactive to both tdTomato and ChAT in the colonic myenteric plexus (Fig. 6C).

**Figure 4:**
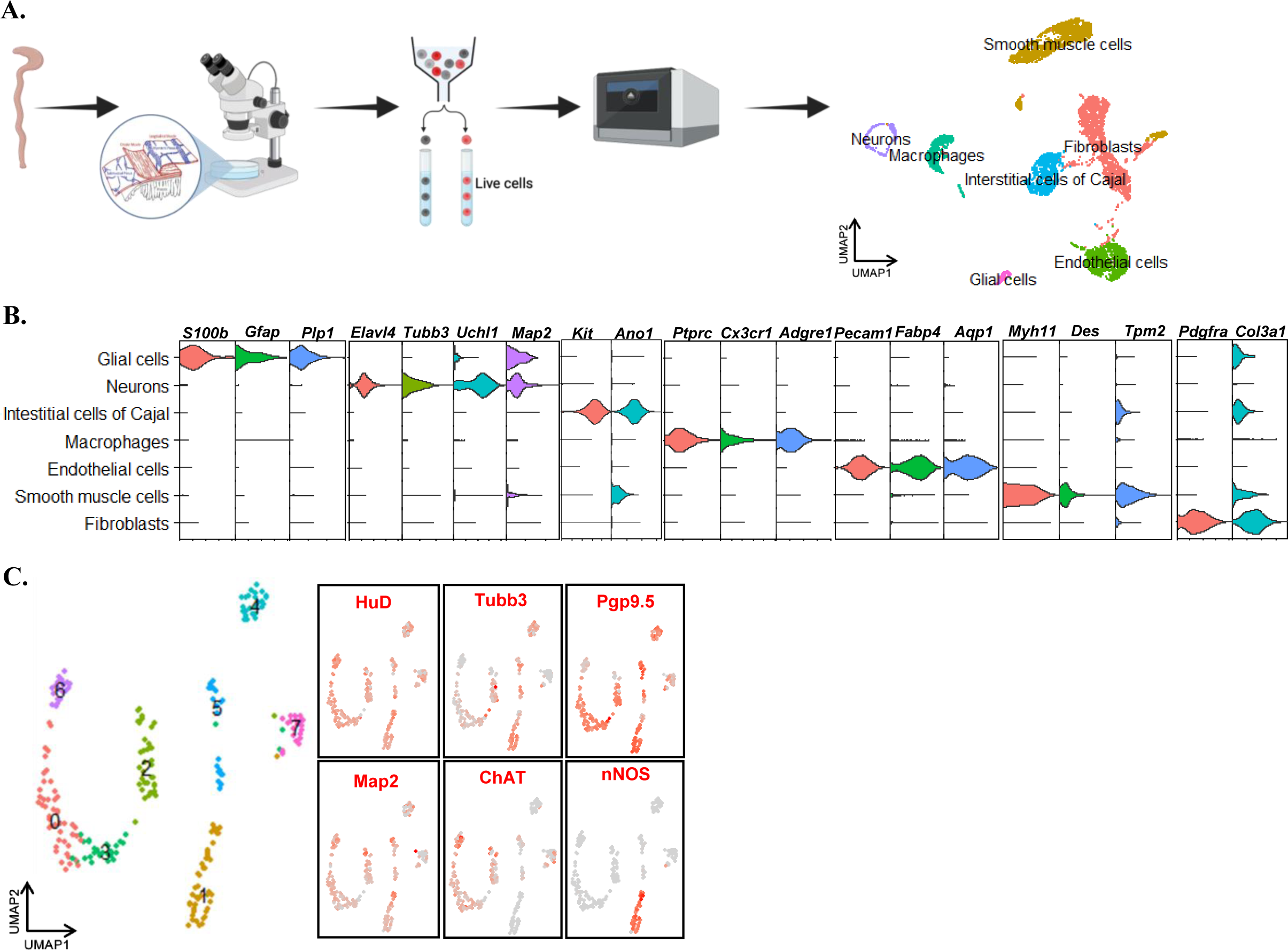
Single-cell RNA sequencing of colonic LMMP cells from healthy adult mice. **A**) Schematic diagram of the colonic longitudinal muscle and myenteric plexus (LMMP) isolation and dissociation for single-cell RNA sequencing (scRNAseq). Live cells were sorted via fluorescence-activated cell sorting (FACS) then gene expression libraries were prepared using 10X Chromium Genomics. Quality control, normalization, clustering and expression analyses of data was done in R using *Seurat* with default settings. Distinct myenteric cells were identified and displayed as a UMAP plot. **B)** Literature-curated genes shown in the violin plot were used to annotate each clusters in the LMMP single-cell dataset. **C)** UMAP plot depicting seven neuronal cell subpopulations identified by confirming the expression of 4 pan-neuronal marker genes in these populations (*Elavl4*/HuD*, Tubb3,* Uchl1/Pgp9.5, *Map2*). The two main enteric subtypes were also identified: *Chat*/ChAT-expressing cholinergic (Clusters 0, 2, 3, 6) and *Nos1*/nNOS-expressing nitrergic (Cluster 1) neurons.

**Figure 5:**
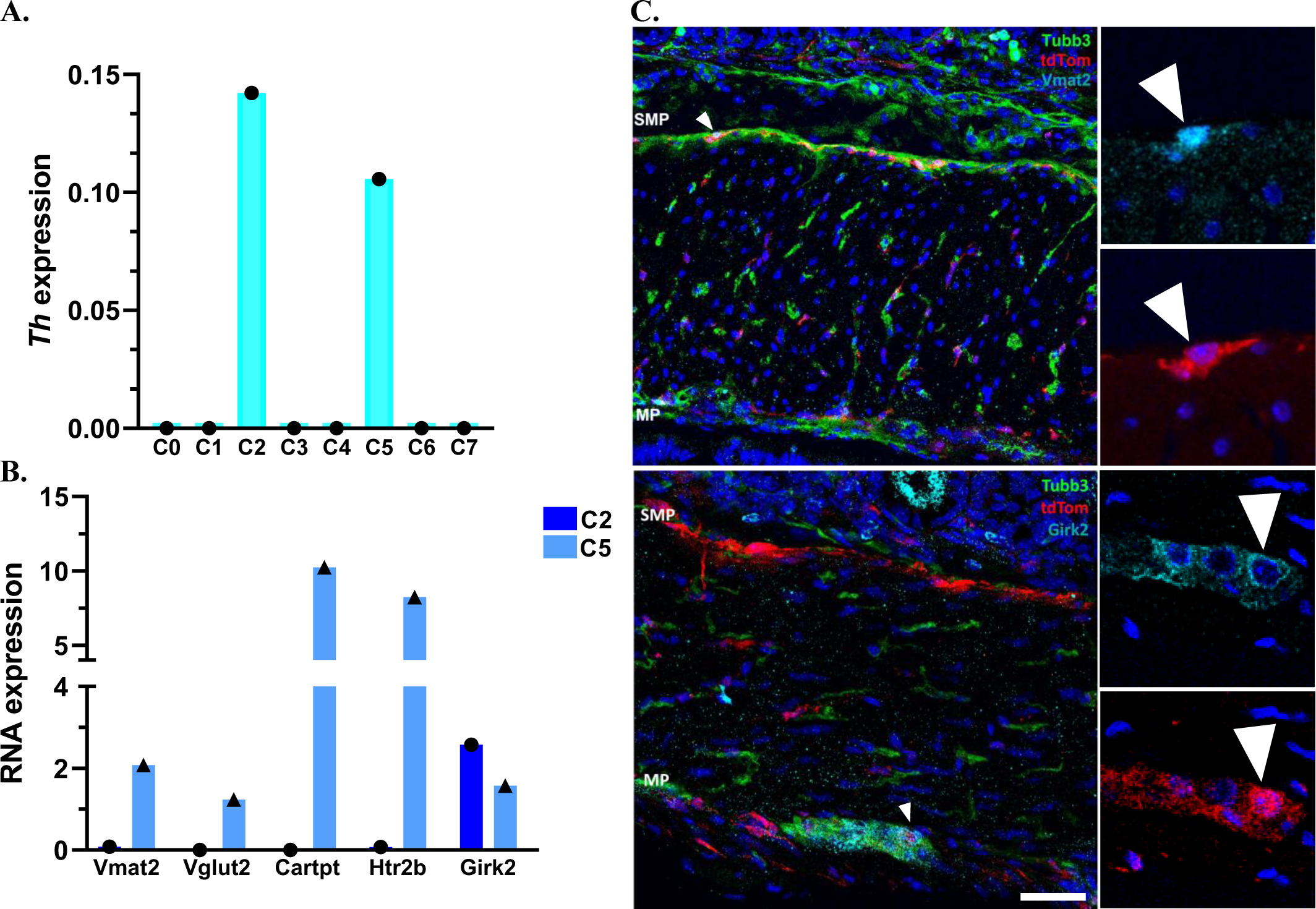
Two dopaminergic neuronal subclusters expressed traditional dopaminergic neuronal markers at varying degrees. **A**) The expression of a canonical DAergic neuronal marker, tyrosine hydroxylase (*Th*), in the neuronal populations are highest in Cluster 2 and 5. **B)** Other previously identified markers of DAergic neurons in the CNS are lower expressed, if any, in Cluster 2 than in 5, suggestive of a potentially distinct *Th*-expressing neuronal populations. **C)** Representative images of colonic tissue demonstrating co-labeling of tdTomato (red) with known dopaminergic neuronal markers (cyan; Vmat2 and Girk2), indicated by arrowheads. Tubb3 is used as pan-neuronal marker (green) and nuclei are stained with Hoechst (blue). Scale bars: 50 μm.

**Figure 6:**
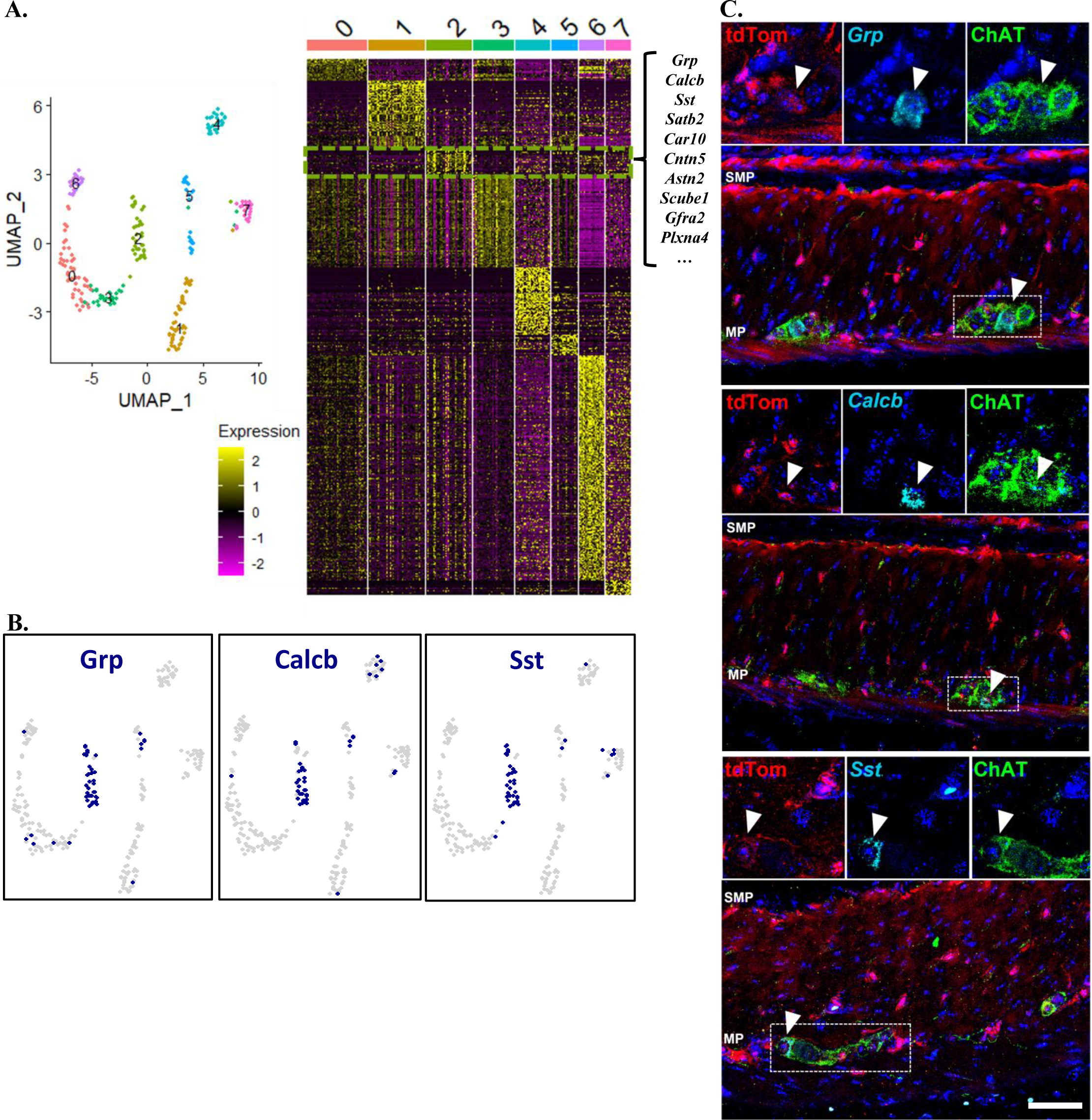
Identification of novel enteric dopaminergic neuron subtype with cholinergic phenotype. **A)** Heatmap indicating upregulated genes (row) in each neuronal cell populations (column). Highlighted in green is DAergic neuronal subtype co-expressing *Th* and *Chat*. **B)** UMAP plots showing novel ENS DA neuron marker genes highly expressed in Cluster 2. **C)** Representative images of colonic tissue demonstrating expression of putative novel DA markers (green; *Grp*, *Calcb* and *Sst*) in the myenteric plexus co-labelled with tdTomato (red) and ChAT (green). Nuclei are stained with Hoechst (blue). Scale bars: 50 μm.

## Discussion

Enteric neurons are numerous and exhibit diverse neuronal phenotypes analogous to the CNS, yet they are largely understudied. DA plays pivotal roles in the brain and dysfunction of DA signaling underpins a plethora of neuropsychiatric disorders and neurodegenerative diseases, including autism, depression and PD (Björklund & Dunnett, 2007; Hamamah et al., 2022). However, the functions of DA neurons in the gut are much less understood with some studies implicating them in gastrointestinal motility (Chalazonitis et al., 2020; Li et al., 2011; Li et al., 2006; Walker et al., 2000). Through advances in DA cell classification in the CNS, striking functions have been found to underscore the heterogeneity of these cell populations in healthy and diseased brains. A better understanding of enteric DA neurons and their vulnerability in disease is critical in the face of mounting evidence suggesting roles of the gut in a myriad of brain disorders (Dinan & Cryan, 2017).

The heterogeneity of DAergic neuronal populations across the plexuses in the gut may account for disparate roles of DA within the ENS. In particular, our discovery of DA neurons with dual-transmitter properties in the gut suggests diversity within the enteric plexus that may contribute to functional differences. Previous work has indeed suggested that enteric DA neurons have multiple roles. For example, deletion of *Dat* in mice suggested the involvement of DAergic neurons in modulating the propagation of colonic migrating motor complexes (CMMC) to promote peristalsis – a central function of the MP (Walker et al., 2000). Meanwhile, intraperitoneal administration of neurotoxins targeting the DA system resulted not only in the ablation of enteric DA neurons, but also reduced colonic mucus content due to decreased signaling through the DA receptor D5 expressed in goblet cells (Li et al., 2019). Contrary to myenteric DA neurons, the implication of DA in mucosal secretion may be attributed to a subset of cells primarily in submucosal ganglia. Submucosal DAergic neurons are also believed to participate in gastrointestinal motility whereby mice with DA neuronal hypoplasia in the SMP manifested longer gastrointestinal transit time and slower colonic expulsion time (Chalazonitis et al., 2020). These submucosal DAergic neuronal subtypes may differ from those responsible for mucosal secretions. Optogenetic stimulation studies in transgenic mice revealed that CMMC generation is mediated in part by the activation of cholinergic neurons while nitrergic neurons are active during its tonic inhibition (Gould et al., 2019). As such, DA release from DAT-tdTomato-positive neurons co-expressing either ChAT or nNOS could be involved in governing this complex process. This is in accordance with previous speculation that repeated patterns of interconnected neural networks in the ENS are activated during peristalsis (Chalazonitis et al., 2022). To add complexity, studies have shown that DA receptor blockade conveys a suppressive action on colonic motility (Li et al., 2011; Li et al., 2006), which refute the previous work mentioned whereby inhibiting DA reuptake elicits fewer smooth muscle contraction (Walker et al., 2000). Clearly, there is a need for a new classification of enteric DA neuron subtypes not only to define which cells control gut motility but also to unveil other intricate DA-related mechanisms.

Given advances in single-cell transcriptomics for profiling DA neurons in the CNS, we also applied this approach in the ENS to unravel novel cellular markers that can demarcate specific DA neuronal subtypes. We found that traditional markers previously shown at high levels in midbrain DA neurons are expressed in enteric neuronal cluster that are also expressing *Th*, which may represent the subpopulation of enteric neurons with functional similarity to DA neurons in the brain. Interestingly, we identified a distinct *Th*-expressing neuronal cluster that may share similar properties with cholinergic neurons as evidenced by its high *Chat* expression. This corroborates our observations using immunohistochemistry, revealing dopaminergic subtypes with dual-transmitter content. This novel *Th-expressing* enteric neuronal subtype also expresses genes found in disparate classes of CNS DA neurons (such as *Grp*, *Sst*, *Calcb*). Prior evidence for the expression of gastrin-releasing neuropeptide encoded by *Grp* in subpopulations of midbrain DA neurons may substantiate its relevance as a candidate marker for DA neuron subtypes in the gut (Kramer et al., 2018; La Manno et al., 2016; Poulin et al., 2014; Viereckel et al., 2016). In contrast to previous reports of widespread expression of GRP-immunoreactive DA neurons in the midbrain (Kramer et al., 2018), our study suggests rare expression which further underscores the heterogeneity of DA subtypes even across the body. As for *Sst* and *Calcb*, DA neurons in the ventral tegmental area (VTA) co-releasing somatostatin (SST) are implicated in the response to stress (Elina et al., 2023) while calcitonin-gene related peptide (CALCB) expressed in hypothalamic DA neurons regulates pain transmission (Charbit et al., 2009). As with the CNS, the manifestation of the dual-transmitter phenotype in the ENS likely contributes to the multifaceted features of the DAergic system in the gut. Further investigations of the differential gene expression landscape of DA neuron subtypes between the ENS and CNS would help better extrapolate shared and unique transcriptional profiles of DA neuron subpopulations and their discrete functions.

It is noteworthy to bear in mind that despite the wealth of information single-cell transcriptomics can now provide, classification of neurons based solely on molecular signatures is likely not sufficient, and thus their morphology, physiological properties and functional connectivity must also be assessed (Zeng & Sanes, 2017). In fact, our data revealed that a small subset of DAT-tdTomato-positive neurons do not express TH, the rate-limiting enzyme for DA synthesis. The reason for this is unclear, albeit between embryonic days 10 to 13, a population of neuronal precursor cells termed transiently catecholaminergic (TC) cells have been reported to display high expression of TH, which progressively diminishes during terminal differentiation (Baetge & Gershon, 1989; Gershon et al., 1984). Although the majority of TC cells give rise to non-catecholaminergic cell types, the question of whether some TC cells express DAT despite low levels of TH is open. Like us, Li and colleagues have indeed demonstrated only partial co-localization of DAT and TH immunoreactivities in the mouse ileum and colon, further alluding to the existence of DAT-expressing neurons that either have no or little expression of TH protein, most prominent in the SMP (Li et al., 2004). It is tempting to speculate that these divergent neuronal subtypes have corresponding functional differences whereby TH neurons may represent a more mature counterpart able to synthesize and release DA while DAT neurons that are TH and VMAT2-negative represent cells that are non-traditional DAergic in the sense that they are not (yet) able to synthesize and release DA.

It remains possible that the DAergic phenotype of tdTomato neurons not expressing TH could be reactivated later in life in response to signals associated with cellular stress. Indeed, in previous work, denervation of axonal projections to the gut culminated in elevated proportions of TH-immunoreactive neurons without affecting the overall number of enteric neurons (Li et al., 2004), implying that existing pools of neurons may upregulate TH expression as a compensatory mechanism in response to injury. Hence, we postulate that a subset of molecularly defined neurons may be acting as *reserves* with submaximal DAergic function. This further begets the question whether the localization of these subtypes (being mostly in SMP) relates to the potential vulnerability of submucosal neurons (given the proximity to the intestinal lumen) and thus presumably necessitates the need to have a pool of *reserve* DA neurons in this region. In agreement with a study depicting the blood-ganglia barrier only at the myenteric plexus (Dora et al., 2021), we also noted the absence of immunoreactivity for Agrin in the SMP, thereby suggesting a lack of protection from the blood-ganglia barrier in the SMP which could render submucosal neurons more susceptible to immune cell and bacterial-related damage. Collectively, these hypotheses can be addressed by marrying detailed knowledge on gene expression patterns in specific DA subtypes with disease or injury models.

A perplexing idea dominating the field of PD is the question of the origin of selective vulnerability of DA neuron subtypes and how specific gene signatures may help us infer the underlying mechanisms (Giguère et al., 2018; Poulin et al., 2020). Although enteric neurodegeneration has been examined in PD, there still exists discrepancies as to the extent and specificity of neuronal damage involved (Lebouvier et al., 2010; McQuade et al., 2021; O’Day et al., 2022; Ohlsson & Englund, 2019). In this regard, the ion channel GIRK2 implicated in modulating neuronal excitability has been hypothesized as a relevant factor that may influence sensitivity of specific DA neurons to degeneration in the CNS. Strong GIRK2 expression is evident in most SNc DAergic cell types, but fewer (with lower levels) in less affected areas, such as the VTA, as described in post-mortem tissues and recent single-cell RNAseq dataset from a murine model of PD (Behzad Yaghmaeian et al., 2023; Reyes et al., 2012). Interestingly, we observed that *Th*-expressing Clusters 2 and 5 express *Girk2*, raising the question of whether comparable selective vulnerability may be present in these gut DA subtypes.

Taken together, our study provides a steppingstone to further garner interest in exploring the diverse molecular and cellular properties of enteric DA neurons. These data suggests that the DAergic neuronal populations in the gut are more heterogenous than previously thought, supporting the notion that DA neurons play diverse functions in the gut. In particular, we identified a novel ChAT-positive dopaminergic neuronal subtype in the gut that co-expresses midbrain DA-related genes (*Grp, Calcb* and *Sst*), exhibiting a dual-transmitter neuronal phenotype.

## Ethics Statement

The animal study was reviewed and approved by the Université de Montréal Animal Care and Use Committee (CDEA).

## Author Contributions

SJR carried out the tissue processing, single-cell preparations and bioinformatic work while SP performed immunostaining, imaging and quantification. SM bred and maintained animal colony. AA created a website allowing access to and *user-friendly* analysis of the enteric neuron single-cell RNAseq data. AM and MY supported tissue processing and bioinformatics, respectively. SG and LET mentored, provided feedback on the project, and contributed to writing the manuscript. SJR and SP prepared figures and drafted the manuscript. SJR and JS conceived the project, supervised the work, and prepared and approved the final manuscript. All authors contributed to the article and approved the submitted version.

## Data availability

Raw and processed single cell data available in GEO (GSE237170) and source code can be found at https://github.com/stratton-lab/recinto-2023-gut_dat_neurons. Raw tabular data of confocal microscopic images available at Zenodo.org: <u>10.5281/zenodo.8231677.</u> Enteric neurons single-cell RNAseq data accessible through a user-interface website available for separate analyses (singlocell.openscience.mcgill.ca/EntericNeurons).

## Acknowledgements

We like to acknowledge Elia Afanasiev for preparing the open-source code available in Github. The study is funded by the joint efforts of The Michael J. Fox Foundation for Parkinson’s Research (MJFF) and the Aligning Science Across Parkinson’s (ASAP) initiative. MJFF administers the grant ASAP 000525 on behalf of ASAP and itself. SJR received a CIHR-Vanier Canada Graduate Scholarship.

## Abbreviations

*Calcb*: Calcitonin-gene related peptide
CNS: central nervous system
ChAT: choline acetyltransferase
*Cartpt*: cocaine- and amphetamine-regulated transcript protein
CMMC: colonic migrating motor complexes
DA: dopamine
DAT: dopamine transporter
ENS: enteric nervous system
FACS: fluorescence-activated cell sorting
*Grp*: gastrin-releasing neuropeptide
*Girk2*: G-protein-regulated inward-rectifier potassium 2
*Htr2b*: 5-hydroxytryptamine/serotonin receptor 2B
LMMP: longitudinal muscle myenteric plexus
MP: myenteric plexus
nNOS: neuronal-specific nitric oxide synthase
PFA: paraformaldehyde
PD: Parkinson’s disease
PNS: peripheral nervous system
RNAseq: RNAsequencing
*Sst*: somatostatin
SMP: submucosal plexus
SNc: substantia nigra pars compacta
TH: tyrosine hydroxylase
VTA: ventral tegmental area
*Vglut2*: vesicular glutamate transporter 2
*Vmat2*: vesicular monoamine transporter 2

